# scHDeepInsight: A Hierarchical Deep Learning Framework for Precise Immune Cell Annotation in Single-Cell RNA-seq Data

**DOI:** 10.1101/2025.06.23.661045

**Authors:** Shangru Jia, Artem Lysenko, Keith A Boroevich, Alok Sharma, Tatsuhiko Tsunoda

## Abstract

Immune cell classification from single-cell RNA sequencing (scRNA-seq) presents significant challenges due to complex hierarchical relationships among cell types. We introduce scHDeepInsight, a deep learning framework that extends our previous scDeepInsight model by integrating a biologically-informed classification architecture with an adaptive hierarchical focal loss. The framework leverages our established method of transforming gene expression data into two-dimensional structured images for use with convolutional neural networks by effectively capturing both global and fine-grained transcriptomic features, overcoming the limitations of flat classification approaches that ignore hierarchical relationships between cell types. scHDeepInsight dynamically adjusts loss contributions to balance performance across the hierarchy levels. It also employs STACAS batch correction, robust random masking, and SHAP-based interpretability to enhance prediction accuracy and biological insight. Comprehensive benchmarking across seven diverse tissue datasets shows scHDeepInsight achieves an average accuracy of 93.2%, representing a 5.1 percentage point improvement over current state-of-the-art methods. The model successfully distinguishes 50 distinct immune cell subtypes with high accuracy, demonstrating proficiency for identifying rare and closely related cell subtypes. These advantages make scHDeepInsight a robust tool for high-resolution immune cell subtype characterization, well suited for detailed immune profiling in immunological studies.

## 1. Introduction

The advent of single-cell RNA sequencing (scRNA-seq) technology has revolutionized our ability to analyze gene expression profiles at an unprecedented resolution, enabling detailed characterization of cellular heterogeneity at the individual cell level. Although existing annotation tools achieve high accuracy in broad cell type identification, they typically do not incorporate predefined immune cell hierarchies during annotation, thus limiting their utility for detailed subtype classification.

Methods such as SingleR [1], Azimuth [2], and scmap [3] are widely used for cell type annotation. These approaches are reference-based, meaning that they typically rely on the similarity metrics calculated between query cells and reference profiles to assign cell type labels. However, these comparisons are generally processed in a uniform manner, disregarding the hierarchical relationships among cell types. Such flat classification strategies can struggle to distinguish closely related subtypes, which often share high transcriptional similarity but differ significantly in function and biological roles. As a result, downstream analyses may suffer from reduced biological relevance and interpretability.

Integrating a hierarchical structure into the annotation process potentially addresses these limitations by explicitly modeling the known biological hierarchy of immune cell populations. Such hierarchical modeling naturally captures functional relationships and widely accepted immune lineage structures, which are partially aligned with known differentiation patterns but are derived from curated ontological annotations, thereby enhancing the biological accuracy and context-specific interpretation of annotation results.

To realize these important improvements, we introduce scHDeepInsight, an enhanced deep learning framework built upon the foundation of scDeepInsight [4] and explicitly incorporates a hierarchical annotation architecture. This design allows the model to better capture nuanced differences between immune cell subtypes, substantially improving the resolution and accuracy of cell type annotations compared to existing methods. Core innovation of scHDeepInsight lies in the integration of three key components: (1) the transformation of high-dimensional gene expression data into spatially-organized 2D images [5] suitable for convolutional neural network (CNN) [6] processing, (2) the implementation of a multi-level classification architecture that preserves biological hierarchies of base-types and subtypes of cells, and (3) the incorporation of an adaptive hierarchical focal loss function that automatically balances training priorities by adjusting the weights of base-type and subtype focal losses according to their relative performance. Through this approach, researchers can precisely dissect the complexities of the immune system at a higher resolution than was previously possible with existing tools. The multi-level classification structure further enables stratified feature importance quantification through SHAP (SHapley Additive exPlanations) [7] analysis at both base-type and subtype cell levels, providing biological insights into the distinct gene expression signatures that differentiate closely related immune cell populations.

In summary, scHDeepInsight provides a robust annotation framework that addresses the limitations of traditional approaches by explicitly modeling hierarchical cell type relationships. This method offers a scientifically rigorous solution for immune cell classification, yielding both improved accuracy and enhanced biological interpretability. As single-cell technologies evolve, this hierarchical classification approach will contribute to advancing our ability to dissect immune cell subpopulations and their functional roles in sophisticated biological processes and disease mechanisms.

## 2. Methods

### 2.1 Overview of scHDeepInsight

scHDeepInsight is a computational framework for single-cell data that enables hierarchical immune cell annotation, rare population detection, and biological interpretation through SHAP analysis. It applies a structured multi-level classification process that reflects known immune cell lineage relationships and supports fine-grained subtype resolution. The workflow begins with data preprocessing and batch-effect correction using STACAS [8], followed by the transformation of gene expression profiles into two-dimensional images and gene selection (via Scanpy [9] to retain the top 5,000 highly expressed genes), and finally hierarchical cell type classification by a convolutional neural network. The last step is done with an automated cell classification model trained on a comprehensive reference atlas composed of immune cell scRNA-seq profiles collected from diverse tissues.

As illustrated in Figure 1, the workflow consists of two main stages: (A) Training phase - Gene expression vectors are transformed into 2D images via DeepInsight followed by random masking, and then processed by a CNN feature extractor and the hierarchical classification architecture with integrated loss function for model training. (B) Application phase - scHDeepInsight performs batch effect correction on query datasets by transforming them using the analogous image conversion procedure followed by the hierarchically-trained CNN to predict both primary cell types and their subtypes. This hierarchical classification strategy enhances both the accuracy of cell type prediction and the biological coherence of the results, particularly in identifying rare immune cell subpopulations and resolving closely related subtypes.

**Figure 1:**
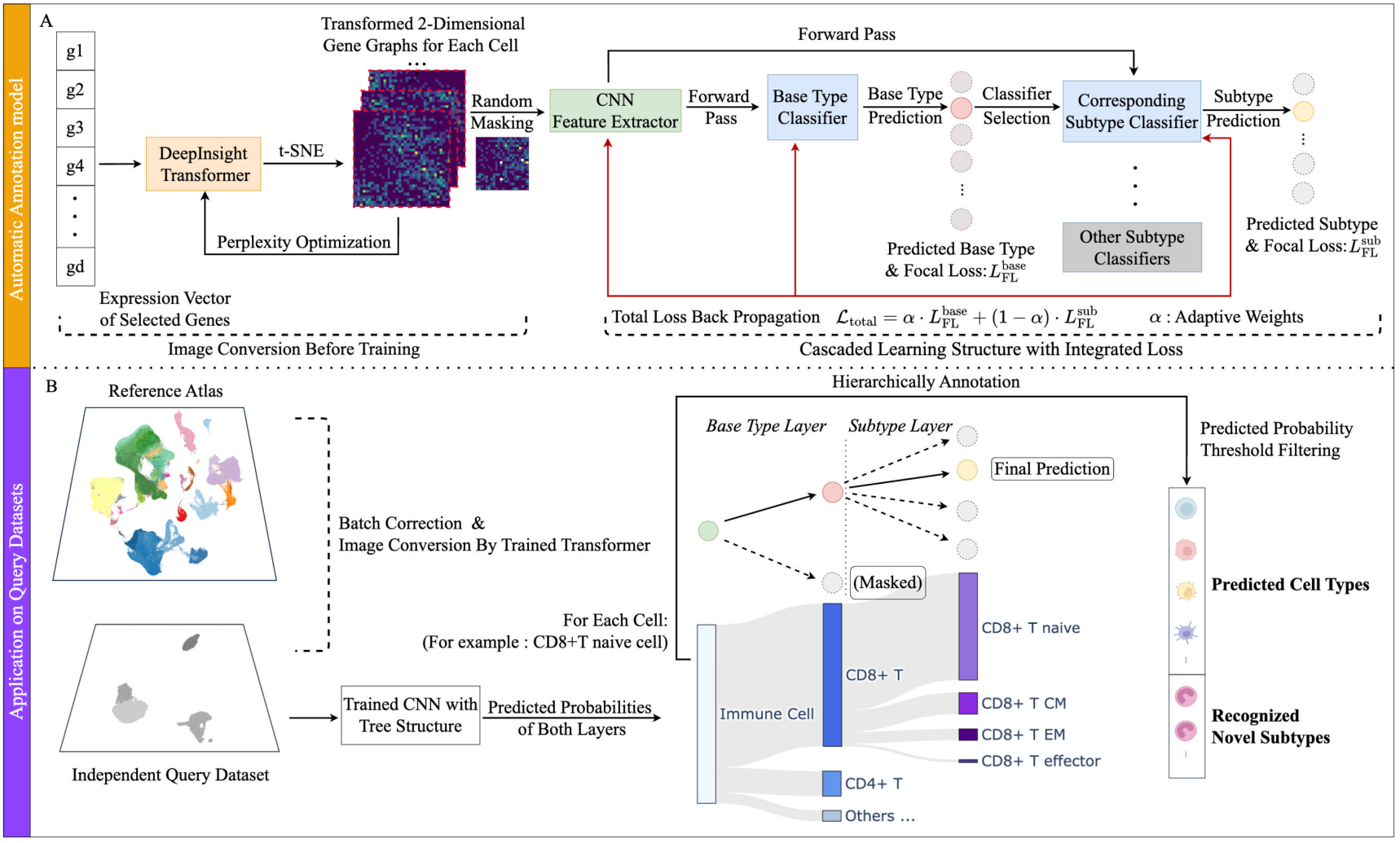
Overview of the scHDeepInsight framework. **(A)** Each cell’s gene expression vector is transformed into a 2D image via DeepInsight [5] (t-SNE [10] with perplexity optimization), optionally masked for data augmentation, then fed into an EfficientNet-B5 [11] CNN trained with a multi-level loss to classify cells hierarchically. **(B)** For query data, after batch correction and image conversion, the trained CNN outputs base-type and subtype probabilities. Masking excludes irrelevant subtypes, enabling hierarchical annotation and uncovering potentially new immune populations.

During the training phase, scHDeepInsight transforms preprocessed gene expression profiles into two-dimensional gene expression images constructed from a reference atlas. These images are then used to train a convolutional neural network based on the EfficientNet-B5 architecture. Unlike the original scDeepInsight, which employs a flat classification approach, scHDeepInsight integrates a multi-level loss function that specifically preserves and reinforces the hierarchical relationships between cell types during classification. This architectural innovation enables a two-stage classification process: first identifying the primary immune cell types, then further refining subtype classification within each lineage.

In the application (test) phase, scHDeepInsight is applied to independent query datasets. The batch-corrected datasets are then fed into the pre-trained CNN, which outputs predicted probabilities for both primary cell types and subtypes. By leveraging the hierarchical structure of predicted probabilities, scHDeepInsight enhances the classification of potentially novel immune cell subpopulations. For rare cell identification, scHDeepInsight employs a probability-based detection mechanism that analyzes the discrepancy patterns between base-type and subtype prediction confidences.

### 2.2 Data Collection

The scHDeepInsight framework was developed and validated using scRNA-seq data from ten published studies [2,12–20], covering a wide range of tissues (e.g., blood, lung, intestine), as detailed in Supplementary_Table 1. In total, the reference dataset includes over 460,000 cells from healthy donors, providing a robust baseline for investigating immune cell heterogeneity. After rigorous preprocessing and STACAS-based batch correction (as described in the Supplementary_Note), we constructed an integrated reference atlas centered on the top 5,000 highly variable genes within the data from the reference datasets, encompassing 15 base immune cell types and more than 50 subtypes. The data integration pipeline follows a systematic workflow from initial study selection through quality control, normalization, and batch correction (Figure 2A). The resulting low-dimensional embedding of the reference atlas shows distinct clusters corresponding to different immune cell types and subtypes (Figure 2B). This hierarchical organization preserves immune cell lineage relationships, ensuring clear separation between major lineages while maintaining biological continuity among related subtypes. This hierarchical reference atlas serves as the foundation for scHDeepInsight’s multi-level classification framework, enabling improved annotation of new query cells while maintaining biological meaningful relationships.

**Figure 2:**
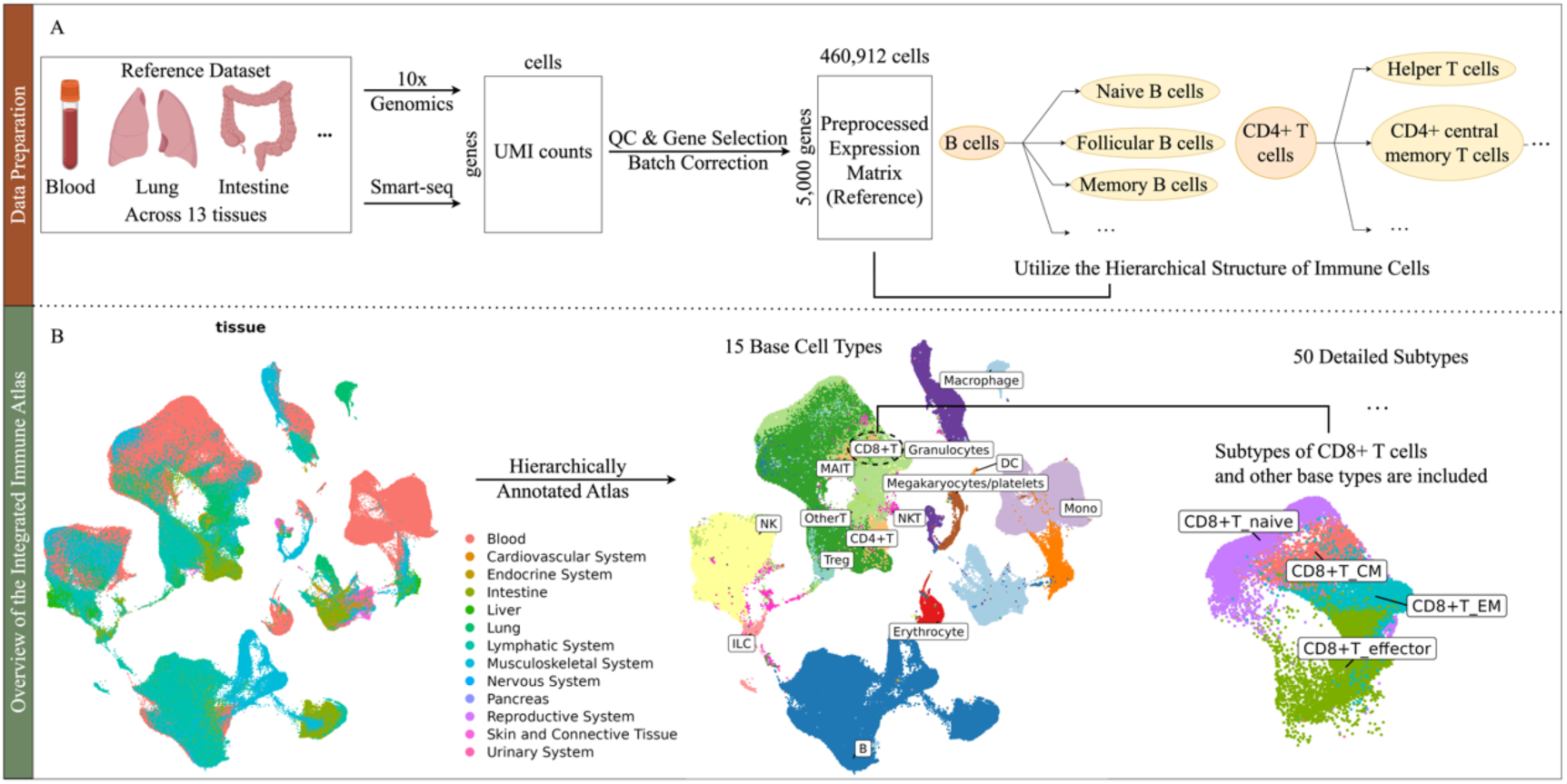
**(A)** Schematic of data integration, from study selection through quality control (QC), normalization, and batch correction. **(B)** Low-dimensional visualization of the final reference atlas comprising 15 base cell types and over 50 subtypes. This hierarchical reference supports scHDeepInsight’s multi-level classification of new (query) cells.

**Table 1:** Summary of datasets used for benchmarking scHDeepInsight and the other tools. Gene coverage percentage indicates the proportion of genes in each query dataset that overlaps with the 5,000 highly variable genes used in the model training.

| Dataset | Tissue | Protocol | Number of cells | Gene Coverage Percentage | Number of celltypes |
| --- | --- | --- | --- | --- | --- |
| Arunachalam et al., 2020 | blood | 10x 3'v2, 10x 3'v3 | 25,954 | 99.8% | 22 |
| Lee et al., 2020 | blood | 10x 3'v3 | 16,298 | 99.5% | 23 |
| Schulte-Schrepping et al., 2020 | blood | 10x 3'v2, 10x 3'v3 | 45,787 | 99.6% | 24 |
| Adams et al. 2020 | lung | 10x 3'v2 | 10,934 | 96.4% | 21 |
| Travaglini et al. 2020 | lung | 10x 3'v3 | 35,699 | 94.6% | 24 |
| MacParland et al. 2018 | liver | 10x 3'v2 | 4,436 | 91.2% | 18 |
| Ledergor et al. 2018 | bone marrow | MARS-seq | 6,367 | 96.3% | 6 |

The query (test) datasets utilized in this study for the evaluation were sourced from various public databases, each covering different tissues and cell types. The datasets include peripheral blood mononuclear cells (PBMCs), lung and liver tissue samples, and bone marrow cells (Table 1). They were generated using a range of scRNA-seq protocols, including multiple versions of 10x Genomics [21] (3’ v2, v3) and MARS-seq [22]. This diversity in datasets and protocols was important for evaluating the adaptability and generalizability of scHDeepInsight across different experimental conditions.

### 2.3 Conversion of Tabular Data into Images

To transform high-dimensional scRNA-seq data into CNN-compatible 2D image representations, pyDeepInsight tool (https://github.com/alok-ai-lab/pyDeepInsight), based on the DeepInsight framework, is employed. In this approach, each gene is mapped to a specific pixel location, and pixel intensity reflects the corresponding gene expression level. Dimensionality reduction techniques, such as t-SNE and UMAP [23], are applied with optimized perplexity settings to project the data into a 2D space, positioning genes with similar expression patterns in close proximity. These coordinates are then converted into pixel positions, creating images in which the intensities represent gene expression patterns.

Considering that query datasets may lack certain genes present in the reference atlas used for training, random masking is introduced to the generated images, thereby injecting controlled noise to enhance robustness against missing gene features in query data. The resulting 2D representations are then utilized by an EfficientNet-B5 CNN for feature extraction and cell type classification. This image-based approach leverages the CNN’s capacity for feature learning while preserving the spatial structure among genes, thereby capturing the underlying gene-gene interaction patterns.

Furthermore, to ensure an optimal gene-to-pixel assignment and eliminate potential collisions where multiple genes would be mapped to the same pixel location, a Linear Sum Assignment (LSA) algorithm was applied [24]. By minimizing the total distance between the high-dimensional feature points and their corresponding pixel centers, this method guarantees a globally optimal gene-to-pixel mapping, ensuring that each gene is uniquely represented in the image space.

### 2.4 Hierarchical prediction model

To overcome the limitations of classification approaches that treat all cell types as independent labels, scHDeepInsight integrates immune lineage structure directly into its classification architecture and training process. This design allows the model to make structured predictions that reflect known biological organization (the complete hierarchical organization of immune cell types and their relationships is visualized in Supplementary_Figure 1), enabling more precise and biologically meaningful cell type identification.

The hierarchical classification in scHDeepInsight follows a structured process where preprocessed scRNA-seq data are first converted into 2D images (224×224×3) using pyDeepInsight. These images undergo random masking as a data augmentation step to enhance model robustness prior to feature extraction by an EfficientNet-B5 CNN, which extracts feature representations shared across classification stages (Figure 3). These features are first fed into a base-type classifier that identifies the broad immune cell category (the detailed classification architecture, including layers and loss functions, is illustrated in Supplementary_Figure 2). Subsequently, based on this primary classification, the appropriate subtype classifier is activated, enabling more refined identification within that cellular lineage. This two-stage approach reflects the biological organization of immune cells.

**Figure 3:**
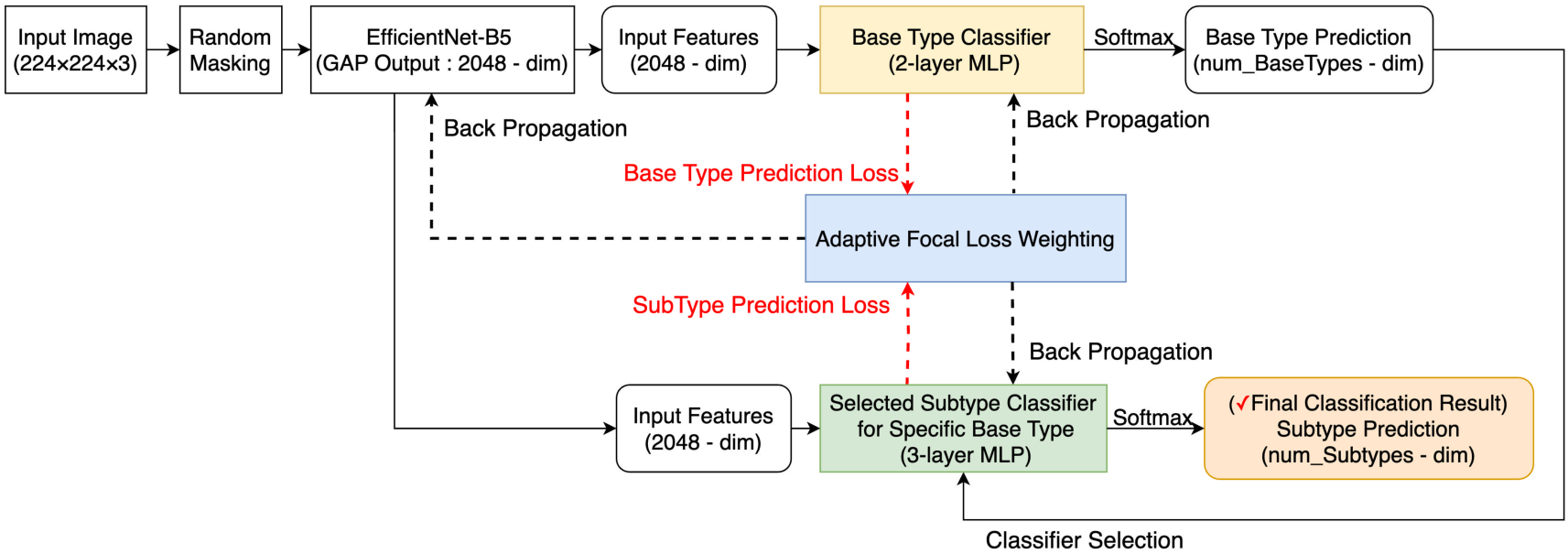
Hierarchical classification model for single-cell type identification. The model uses EfficientNet-B5 to extract features from transformed gene expression images for sequential classification of cell types and subtypes. Focal loss [25] is applied to address the imbalance of different cell types in single-cell data. During training, loss signals back-propagate through the entire network, optimizing both classification levels simultaneously. The adaptive weighting mechanism balances base-type and subtype losses throughout the training process. This hierarchical approach improves discrimination between closely related cell subtypes.

A multi-level loss function optimizes predictions at both base-type and subtype levels simultaneously, with error signals propagated back through the network to capture hierarchical dependencies between cell types. Additionally, probability masking sets the probabilities of subtypes outside the predicted primary category to zero, ensuring that downstream classifiers operate only within biologically relevant subtype spaces and reducing misclassification across unrelated lineages. In cases where a base-type has no defined subtypes, it is treated as a terminal leaf in the classification tree. The model bypasses the subtype classifier for such nodes, and the base-type prediction itself serves as the final output. This implementation preserves hierarchical consistency while accommodating the limited subtype resolution available in current immune reference annotations.

This framework enables the model to learn biologically meaningful cell relationships while leveraging shared feature representations to improve generalization. The following section details the implementation of the multi-level loss function that refines classification performance across hierarchical levels.

### 2.5 Adaptive Hierarchical Focal Loss

For effective hierarchical image classification, the model must address class imbalance while maintaining predictive accuracy across different granularity levels. To achieve this, scHDeepInsight employs an Adaptive Hierarchical Focal Loss (AHFL) that optimizes classification at multiple levels of the hierarchy. The AHFL is formalized in Equations (1) and (2) below:

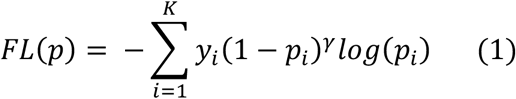

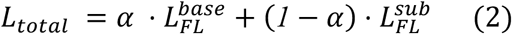

Here, 𝐾 represents the total number of possible classes, 𝑦_1_ ∈ {0,1} denotes the true label for class 𝑖 , and 𝑝_i_ is the predicted probability for that class. The focusing parameter 𝛾 modulates the influence of well-classified versus hard-to-classify examples. Following prior work and to better handle class imbalance in single-cell RNA-seq data, we set 𝛾 = 3.0 used in all experiments. This focusing mechanism helps address the substantial class imbalance frequently observed in single-cell RNA-seq datasets, where certain immune cell subtypes may be underrepresented. By down-weighting the loss contribution from easy examples, the model focuses more on rare or ambiguous populations, improving classification robustness in imbalanced settings. 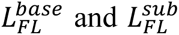 represent the focal losses computed at the base and subtype levels, respectively, and 𝛼 ∈ {0,1} balances their relative contributions. Unlike static weighting schemes, 𝛼 is dynamically updated during training based on the relative difficulty of the two classification levels. As defined in Equation (3), this dynamic adjustment helps the model emphasize the more challenging task at each iteration:

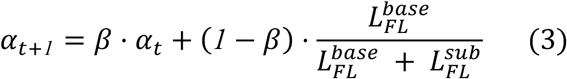

This adaptive weighting mechanism dynamically directs the model’s focus towards the most challenging hierarchical level during training, depending on the relative difficulty of the classification tasks at each level. When base-type classification becomes more accurate 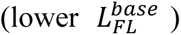, the algorithm shifts focus toward improving subtype classification by decreasing 𝛼*_t_* in the next iteration. Conversely, when subtype classification improves substantially, the model allocates more weight to refining base-type predictions. By dynamically balancing the emphasis between broader classification tasks and finer-grained ones, the model achieves robust and accurate predictions across the entire hierarchy.

Through joint backpropagation, the framework simultaneously optimizes the loss at the base immune cell type level and at the subtype level, ensuring that classification errors at each level are effectively corrected. This approach not only improves overall immune cell type classification accuracy but also enhances the biological interpretability of the results, making them consistent with the known hierarchical structure of immune cells. This is particularly crucial when analyzing complex immune cell data, where scHDeepInsight can accurately capture the hierarchical relationships and functional states of immune cells.

The multi-level loss function design provides robust support for scHDeepInsight, enabling it to deliver precise and detailed immune cell classifications in scRNA-seq data analysis, thereby contributing to a deep understanding of the complexity and diversity of the immune system. The AHFL approach implemented in scHDeepInsight provides a framework for characterizing immune cellular heterogeneity at multiple granularity levels, allowing for more nuanced classification of both common and rare immune cell populations.

## 3. Results

### 3.1 Benchmarking with State-of-the-Art Methods

To evaluate the performance of scHDeepInsight (Methods), a series of benchmarking experiments were conducted using seven independent query datasets derived from diverse tissues. These datasets were selected to represent a diverse array of tissues, cell types, and disease conditions, providing a testbed for assessing the accuracy, precision, and robustness of scHDeepInsight compared to other state-of-the-art (SOTA) cell annotation methods, including SingleR [1], Azimuth [2], scDeepInsight [4], CellTypist [12], Garnett [26], scType [27], and GPTCellType [28]. **The evaluation metrics and technical summaries of these benchmarked methods are provided in the Supplementary_Note**.

#### 3.1.1 Benchmarking Evaluation Across Multiple Metrics

This evaluation was performed at the subtype level classification, which allows assessment of the model’s ability to distinguish between specific immune cell subtypes. scHDeepInsight demonstrated consistently robust performance, achieving an average accuracy of 93.2% and precision of 91.1% across diverse datasets (Figure 4A, 4B; the detailed training and validation accuracy trends are provided in Supplementary_Figure 3). The method achieved an F1-score of 90.5%, reflecting a balance between precision and recall, alongside an AUPRC of 89.7%, a metric particularly suitable for evaluating classification performance on datasets with imbalanced class distributions (Figure 4C, 4D).

**Figure 4:**
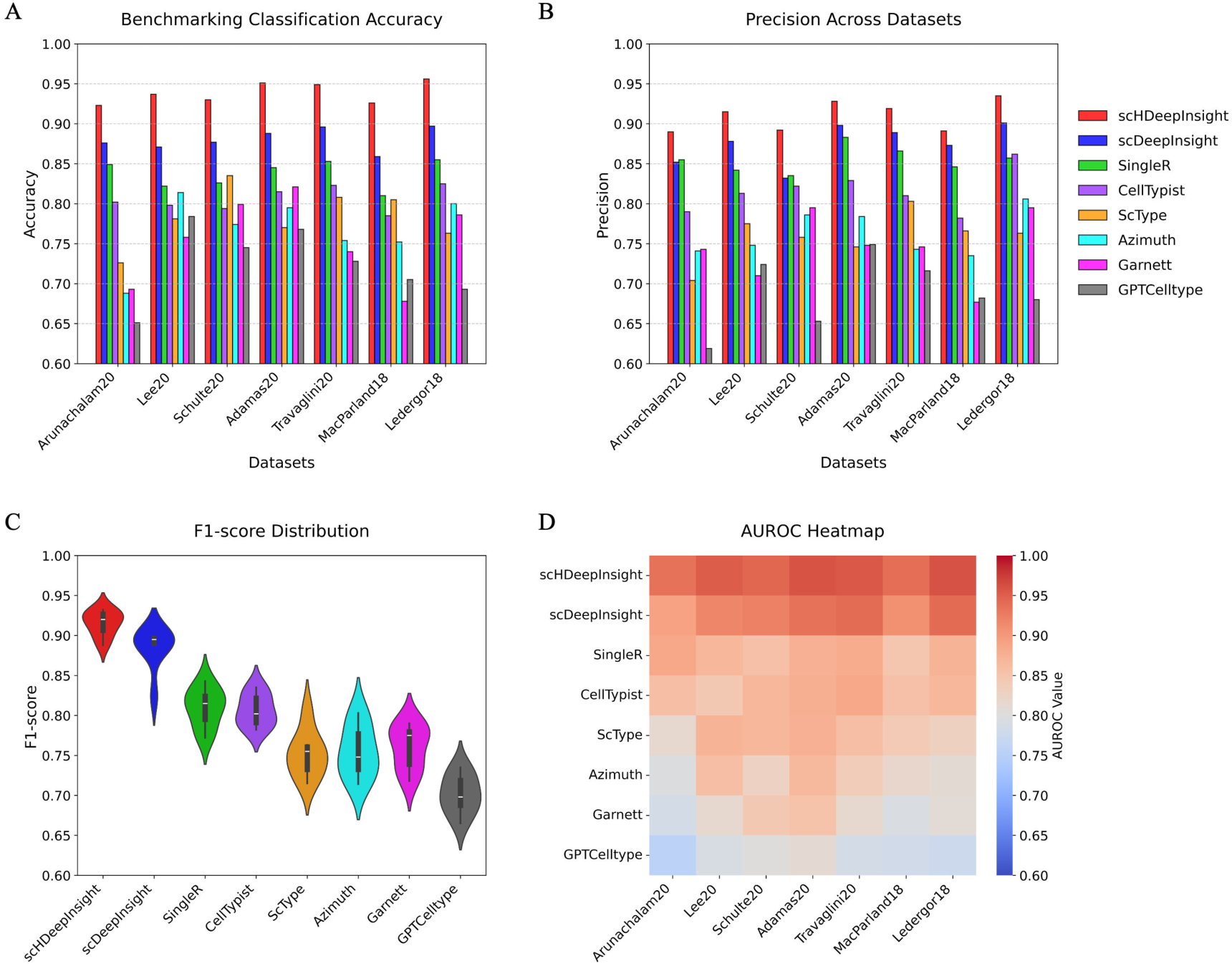
Benchmarking results. **(A)** Accuracies across seven datasets for all classification methods (scHDeepInsight in red). **(B)** Precisions for the same methods. **(C)** F1-score distributions as violin plots revealing median performance and variability across methods. **(D)** AUPRC heatmap displaying classification strength with color intensity corresponding to performance values.

Comparative analysis with scDeepInsight (the next highest performing method) revealed improvements of 5.1% in accuracy, 3.3% in precision, 3.1% in F1-score, and 3.6% in AUPRC (detailed comparative results for all methods are shown in Supplementary_Table 2). These improvements across all evaluation metrics indicate scHDeepInsight’s enhanced classification capability across the diverse cell types and datasets tested, as illustrated in the performance plots and the heatmap.

#### 3.1.2 Fine-grained Immune Cell Classification

Owing to the hierarchical classification design and comprehensive reference atlas covering 50 immune cell subtypes, scHDeepInsight demonstrates enhanced capacity to accurately discriminate between closely related subtypes that are often indistinguishable by conventional annotation methods. For instance, in the PBMC query dataset (Lee) [29], scHDeepInsight successfully distinguished between closely related subtypes, achieving high classification accuracy with minimal misclassification errors (Figure 5A). In the labial gland dataset Pranzatelli [30], scHDeepInsight successfully recovered the distinct immunoglobulin-based subtypes (IgA+ and IgG+) of plasma cells originally labeled by experts (Figure 5B, 5C). However, other supervised annotation methods such as CellTypist failed to maintain this resolution, instead grouping all plasma cells into a single broad category (Figure 5D). By delineating these subtle transcriptional differences, scHDeepInsight highlights the distinct advantage provided by the hierarchical classification framework in accurately resolving closely related immune cell populations that might otherwise be pooled into a single category.

**Figure 5:**
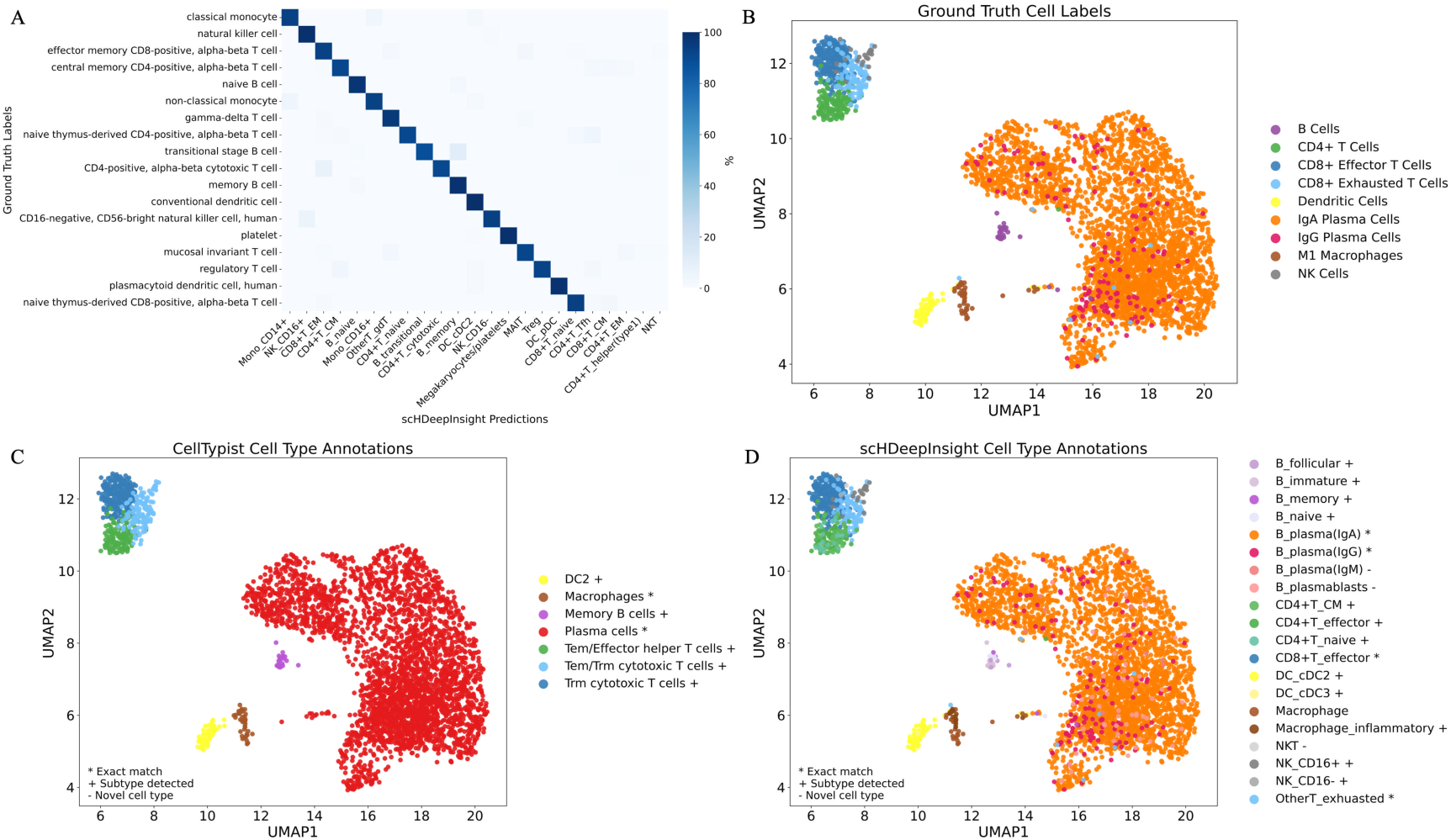
Classification results demonstrating immune cell subtype identification capabilities. **(A)** The confusion matrix of scHDeepInsight predictions on the Lee dataset. **(B)** UMAP visualization of the labial gland dataset with original expert annotations, including detailed plasma cell subtypes. **(C)** scHDeepInsight classifications on the labial gland dataset shows accurate classifications of cell types and subtypes that match with the original annotations, alongside the identification of potentially novel cell populations. **(D)** Annotation results with CellTypist on the labial gland dataset.

#### 3.1.3 Conclusion of the benchmarking

Benchmarking metrics computed at the subtype level across 50 immune cell categories demonstrated that scHDeepInsight achieves high classification accuracy not only for broad immune cell types but also for closely related and biologically challenging subtypes. Furthermore, it successfully resolves biologically meaningful subtypes— such as functionally distinct plasma cell populations—that are often indistinguishable by other annotation methods, highlighting its capacity to provide annotations that are both accurate and biologically interpretable. By combining the hierarchical classification scheme with the robust data preprocessing and the batch correction, scHDeepInsight provides reliable and high-resolution annotations that are well suited for downstream immunological analyses.

### 3.2 Recognition of rare cell types

scHDeepInsight’s hierarchical classification framework enables effective detection of novel cell populations through analysis of prediction probability patterns. The model assigns probability scores at both base and subtype levels, creating a quantitative signature that reflects cellular identity with greater nuance than conventional binary classification approaches (Methods).

In the glioblastoma dataset Ruiz-Moreno [31], immune cells infiltrating the tumor microenvironment demonstrated distinctive probability patterns (Figure 6A). Cells assigned to known cell types in the reference tend to exhibit high prediction probabilities at both the base-type and subtype levels (Figure 6B, 6C), whereas novel populations such as glioma-associated immune cells often display high base-type confidence but reduced confidence at the subtype level. This is exemplified by cells transcriptionally classified as microglial cells, which show high base-type probabilities for the macrophage lineage while lacking strong subtype-level assignment (Figure 6C, 6D).

**Figure 6:**
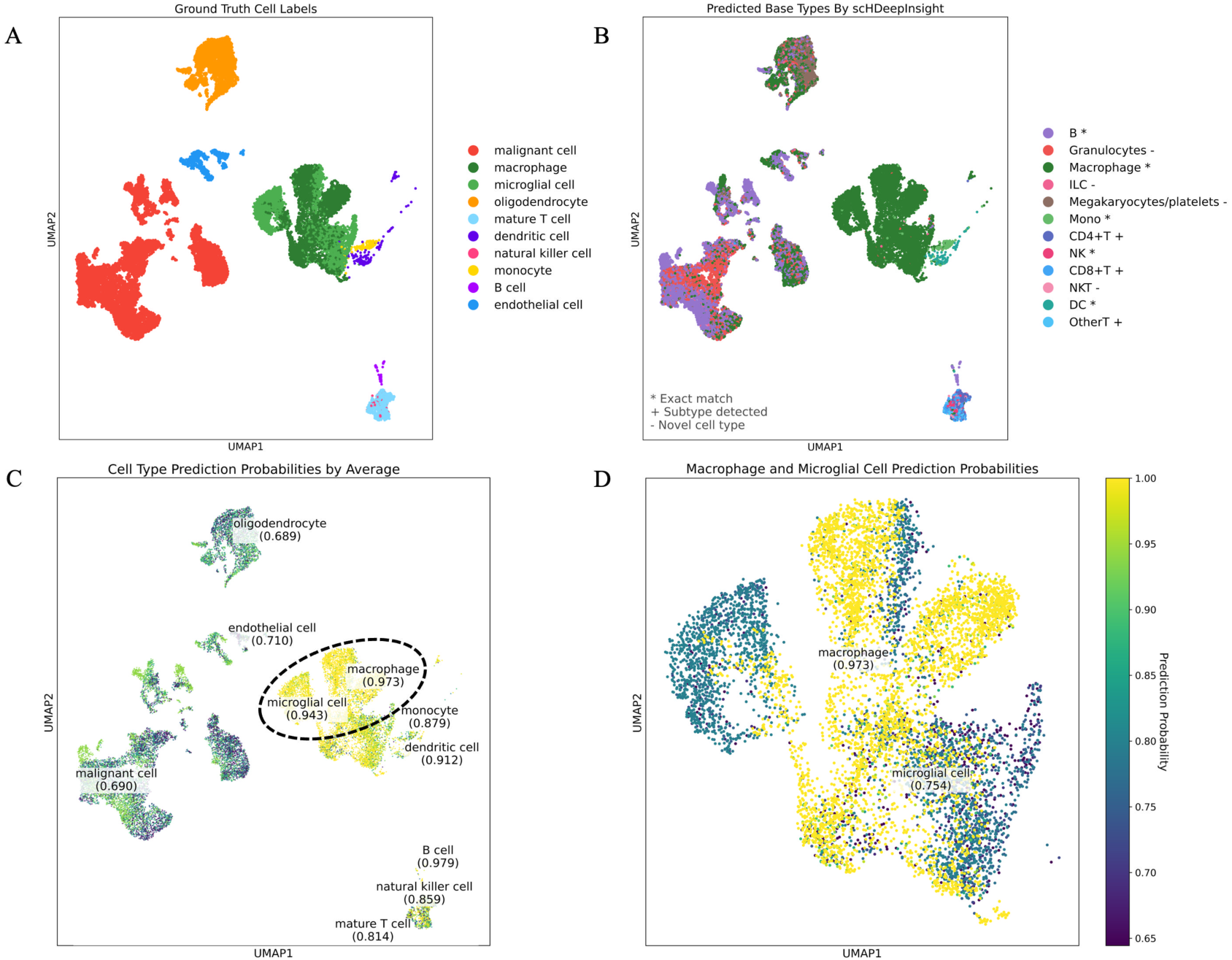
Identification of novel cell populations through hierarchical probability analysis. **(A)** UMAP visualization with ground truth annotations. **(B)** Prediction score distribution by cell type. **(C)** Base-type probabilities on UMAP embedding. **(D)** Relationship between base-type and subtype probabilities revealing signatures of rare cell states.

These findings illustrate that scHDeepInsight’s probabilistic output enables a graded representation of cellular identity, capturing both canonical immune subtypes and cell populations that diverge from transcriptional profiles represented in the reference atlas. The model’s ability to represent prediction uncertainty across hierarchical levels moves beyond discrete label assignment, allowing a more continuous and biologically informative characterization of cell identity. This approach is particularly valuable in heterogeneous contexts such as the tumor microenvironment, where immune populations may exhibit context-specific transcriptional programs and deviate from canonical definitions.

### 3.3 Analysis of SHAP-based Feature Importance for Immune Cell Classification

scHDeepInsight employs SHAP analysis to identify discriminative genomic features critical for accurate cell type classification at both base and subtype levels. This approach quantitatively measures the contribution of each gene to the model’s classification decisions across hierarchical levels. SHAP analysis was conducted separately for each immune cell type and subtype, using 300 samples as its background reference and 1000 samples for the analysis. The resulting values represent the average feature importance across cells within each specific class, thereby enabling identification of both lineage-defining and subtype-specific gene signatures. As shown in Figure 7, the model identifies distinct gene importance patterns across different immune cell populations, with both expected canonical markers and subtype-specific genes revealed as predictive features.

**Figure 7:**
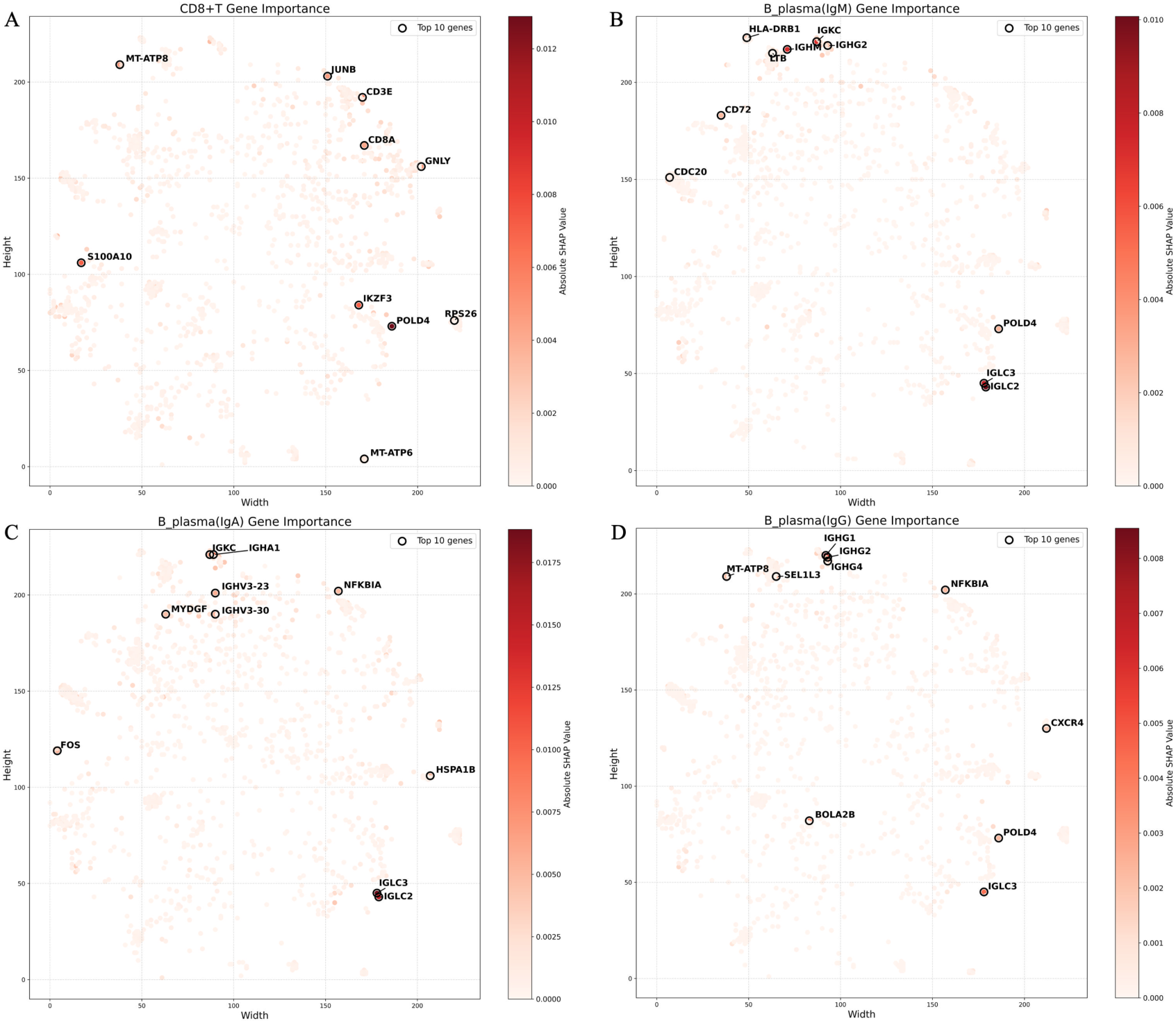
SHAP-based gene importance analysis for immune cell classification. **(A)** CD8+ T cell gene importance, highlighting CD8A, CD3E, and effector molecules like GNLY. **(B)** IgM-expressing plasma cell gene importance, with high values for IGHM. **(C)** IgA-expressing plasma cell gene importance, showing elevated SHAP values for IGHA1 and associated light chain genes. **(D)** IgG-expressing plasma cell gene importance, featuring IGHG1, IGHG2.

For CD8+ T cells (Figure 7A), canonical T cell markers CD8A and CD3E demonstrate high positive SHAP values, confirming their expected role in defining cytotoxic T cell identity. Similarly, the cytotoxic effector molecule GNLY and transcription factors involved in T cell activation contribute positively to classification decisions. Notably, SHAP analysis can capture both positive and negative gene signatures that collectively define cell subtypes. CD72 is typically downregulated during plasma cell differentiation [32], and this biological phenomenon is further validated by the observation of a highly negative SHAP value associated with CD72 (Figure 7B). This finding highlights the capability of scHDeepInsight not only to leverage the presence of specific transcripts but also to recognize the significance of their absence in classification decisions, thereby enabling more accurate cell type identification.

The hierarchical approach also successfully identifies common plasma cell features across subtypes, such as immunoglobulin light chain genes (IGLC2, IGLC3), while still capturing the isotype-specific differences that biologically distinguish these populations. Each plasma cell subtype exhibits a distinctive SHAP pattern dominated by isotype-specific immunoglobulin heavy chain genes: IGHA1 for IgA-expressing plasma cells (Figure 7C), IGHM for IgM-expressing plasma cells (Figure 7B), and IGHG1/IGHG2 for IgG-expressing plasma cells (Figure 7D). The diversity of these patterns further validates that the detailed subtype classifier within the hierarchical classification framework is capable of capturing both shared lineage features and subtype-specific genes, further supporting its ability to discriminate between closely related cellular states from an immunological perspective.

## 4. Discussion

scHDeepInsight advances immune cell annotation in scRNA-seq by implementing a hierarchical classification framework that reflects the biological organization of immune lineages. In contrast to conventional annotation strategies that treat all cell types as independent categories, it leverages shared representations across hierarchical levels, thereby improving the resolution of both major lineages and transcriptionally similar subtypes.

Beyond improvements in predictive performance, scHDeepInsight incorporates several technical innovations to enhance biological interpretability. The adaptive hierarchical focal loss dynamically balances optimization across classification levels, increasing sensitivity to both coarse-grained and fine-grained cell distinctions. The image-based transformation of gene expression profiles enables convolutional neural networks to learn complex expression patterns from inherently non-spatial data. Additionally, the generation of structured probability distributions across the hierarchy supports a continuous and interpretable representation of cellular identity. This probabilistic design facilitates the detection of potentially novel or context-specific immune populations, particularly when cells exhibit high base-type confidence but low subtype-level assignment—as observed in the glioblastoma dataset. Model interpretability is further strengthened by SHAP-based feature analysis, which reveals gene expression patterns associated with both lineage specification and functional divergence. By identifying canonical markers alongside context-specific regulatory features, the model enhances confidence in subtype predictions and provides insights into the gene expression patterns underlying immune heterogeneity.

Despite these substantial advancements, several limitations persist that merit further exploration. Classification accuracy is constrained by the resolution and completeness of the reference atlas, which represents a fundamental limitation that cannot be fully overcome by the hierarchical structure constructed in this model. Furthermore, the transformation of gene expression vectors into image representations introduces computational overhead relative to vector-based methods, though this cost is often justified in applications requiring high-resolution immune profiling.

Future development of scHDeepInsight will focus on extending its generalizability and biological scope. The integration of multi-omics modalities, such as CITE-seq [33] and spatial transcriptomics, is expected to improve subtype resolution by incorporating protein-level and spatial context information. Self-supervised learning approaches may enhance feature extraction from unlabeled data, enabling discovery of novel immune states without prior annotation. Incorporating pseudotime inference would allow dynamic modeling of differentiation trajectories, extending the framework beyond static classification. Finally, transfer learning strategies offer a promising path toward improving adaptability across tissues or species with minimal retraining, broadening the applicability of scHDeepInsight to diverse biological contexts. As a further extension, incorporating non-immune cell populations into the hierarchical framework would enable unified modeling of diverse cellular systems beyond the immune compartment.

## 5. Conclusion

scHDeepInsight is a deep learning framework for hierarchical immune cell classification using scRNA-seq data, incorporating CNN architectures and immune lineage structures to enhance accuracy and resolution of cell type annotation.

Through the integrated use of the multi-level loss function and adaptive weighting approach, scHDeepInsight achieves accurate identification of both common immune lineages and closely related subtypes while maintaining their hierarchical relationships. The integration of batch effect correction ensures consistent performance across datasets under diverse experimental conditions. The comprehensive benchmarking demonstrated that scHDeepInsight outperformed the existing annotation methods across the multiple performance metrics, particularly in distinguishing the closely related immune cell subtypes within complex tissues. In addition, scHDeepInsight revealed novel immune populations through hierarchical analysis on prediction probabilities, highlighting its potential for uncovering biologically relevant cellular diversity beyond canonical annotations. This improved resolution enables precise characterizations of cellular heterogeneities in immunological research.

As the single-cell technologies continue to evolve, the hierarchical classification approach implemented in scHDeepInsight will be increasingly valuable for advancing our understanding of immune cell diversity and functional specialization.

## Data Availability

All datasets used in this study were obtained from publicly available repositories. The Lee (GSE149689) [29], Pranzatelli (phs002446) [30], Ruiz-Moreno (GSE211376) [31], Adamas [34] (GSE134692), Travaglini [35] (EGAS00001004344), Arunachalam (GSE155673) [36], Schulte (EGAS00001004571) [37], MacParland (GSE115469) [38] and Ledergor (GSE117156) [39] are also accessible via the CELLxGENE [40] portal: https://cellxgene.cziscience.com/datasets. The integrated reference dataset used for model training is available at Figshare (https://doi.org/10.6084/m9.figshare.28831010).

## Code Availability

The source code is publicly available at the GitHub repository: https://github.com/shangruJia/scHDeepInsight. A packaged Python library is also accessible via PyPI (https://pypi.org/project/SCHdeepinsight/) for straightforward installation and use.

## Acknowledgements

The results shown in this paper are in part based upon publicly available single-cell datasets from Gene Expression Omnibus (GEO), the European Genome-phenome Archive (EGA), and the CELLxGENE portal. We appreciate the researchers and consortia responsible for generating and openly sharing these valuable datasets, enabling comprehensive benchmarking and validation performed in this study.

## Author contributions

SJ implemented the whole pipeline, evaluated the performance, and wrote the first draft and contributed to the subsequent versions of the manuscript. AL advised for the model and the evaluation, and contributed to the manuscript writeups. KAB checked the model, and helped in the manuscript writeup. AS and TT perceived, supervised, and contributed to the manuscript writeups. All authors read and approved the manuscript.

